# Orbit Image Analysis: An open-source whole slide image analysis tool

**DOI:** 10.1101/731000

**Authors:** Manuel Stritt, Anna K. Stalder, Enrico Vezzali

## Abstract

We describe the open-source whole slide image analysis tool Orbit Image Analysis. It is a generic tile-processing engine which allows the execution of various image analysis algorithms provided by either Orbit itself or other open-source solutions using a tile-based map-reduce execution framework. We show its sophisticated machine-learning approach for WSI quantification, and its flexibility by integrating a deep learning segmentation method for complex object detection. It can run locally standalone or connect to the open-source image server OMERO, and provides scale-out functionality to use the Spark framework for distributed computing. We demonstrate the application of Orbit in three real-world use-cases: Idiopathic lung fibrosis, nerve fibre density quantification, and glomeruli detection in kidney.

**Author summary:** Whole slide images (WSI) are digital scans of samples, e.g. tissue sections. It is very convenient to view samples in this digital form, and with the increasing computation power it can also be used for quantification. These images are often too large to be analysed with standard tools. To overcome this issue, we created on open-source tool called Orbit Image Analysis which divides the images into smaller parts and allows the analysis of it with either embedded algorithms or the integration of existing tools. It also provides mechanisms to process huge amounts of images in distributed computing environments such as clusters or cloud infrastructures. In this paper we describe the Orbit system and demonstrate its application based on three real-word use-cases.

## Introduction

The use of digital pathology (DP) as a companion diagnostic is growing rapidly, with several reports of pathology departments transitioning to signing out cases using either partial or in some cases completely digital workflows ((1), (2), (3), (4)). This trend, along with the arrival of commercial whole slide imaging (WSI) scanners has helped to drive the adoption of DP as both a research and clinical tool. However, the purpose of acquiring WSI digital data is not just to have a digital, computer-aided display of pathology specimens, but to make it possible to apply advanced computer vision and machine learning tools. Ideally, this will accelerate the processing of slides and cases in research, drug development, clinical trials and clinical diagnosis - in all the domains where pathology makes a critical contribution to understanding the effect of experimental perturbation or disease. A rich set of algorithms using conventional and deep learning tools are now available in either commercial or open source form and the community developing these tools has matured so that community challenges that compare different approaches are now being run and some cases completed ((5), (6)). This maturation has made it possible to build user-facing software platforms that deliver these advanced algorithms in usable, user-directed programs from either commercial developers or open source developers. The first generation of these algorithms, which used intensity-based pixel classification, has been done to quantify those images. While this worked well for simple analysis tasks, for slightly more complex tasks such as lung fibrosis quantification, where structural collagen has the same staining colour as new collagen, they fail completely, whereas for pathologists these tasks seem to be relatively easy. The rationale behind this discrepancy is that humans are able to take the context into consideration, e.g. to understand that a cell lays inside a particular region of a tissue and thus is different from another cell, even if both have the same colour. Simple intensity-based pixel analysis cannot deliver an acceptable result in this case. Instead a more holistic approach is required that combines intensity and colour with more sophisticated features that quantify the context of any object, and include this context in the classification in order to successfully classify samples of lung fibrosis or other challenging tissues. Many feature-based approaches at object- or cell-level have been published to quantify specific use-cases ((7), (8), (9)). With the emergence of deep learning methods even more complex scenarios could be quantified ((10), (11), (12), (13), (14)). To harmonize these approaches and create a generic and modular framework for sophisticated WSI quantification, we developed Orbit Image Analysis - a framework which uses machine learning to understand the context within huge WSI, makes use of this knowledge to quantify structures on different magnification levels, and allows the integration of arbitrary analysis algorithms including deep learning. Orbit is open source software, with the source code and binary executables freely available (www.orbit.bio; https://github.com/mstritt/), licenced under GPLv3.

### Design and implementation

Orbit’s context-based structure classification is based on the so-called structure-size, a surrounding area for each pixel, which is used to compute several features on multiple image resolutions. These features describe the structure of the underlying tissue or other biological sample and are used as an input for a Support Vector Machine (SVM) to discriminate regions within the image. In contrast to deep learning methods where a huge training set is needed, this approach allows the users to specify just a few training regions on-the-fly and create a model within minutes.

In addition to this basic functionality sophisticated image analysis algorithms, such as segmentation and deep learning classification exist in the open-source world, but these algorithms usually work on small, in memory fitting, images and not on very large images. This is important because WSIs are typically quite large, e.g., 150,000 x 60,000 pixels and recent advances in staining and data acquisition produce multiplexed images of these dimensions in 40 different channels or more (15). Orbit’s generic map-reduce based tile-processing approach enables the application of these algorithms for large WSI.

The combination of the context-based structure classification together with the generic tile-processing engine which enables the integration of further algorithms unveils a comprehensive, scriptable and scalable framework for objective WSI quantification.

Another powerful WSI tool is QuPath (16) which focus is on object segmentation in combination with a object-based data structure which allows further analysis on the segmented objects. It has sophisticated data analysis and visualization methods with main target on bright field images. QuPath provides a very nice tool chain starting from Tissue Micro Array (TMA) processing till the result visualization. In contrast to this, Orbit’s main focus is on enabling existing image analysis tools for WSI and thus provides a generic tile-processing engine so that algorithms can be easily integrated into the system. In addition, it comes with sophisticated build-in tools ready to use within a user-friendly interface. A batch mode allows the analysis of hundreds of images in parallel on a scale-out infrastructure, e.g. a Spark cluster which allows image processing on an on-premise cluster or using cloud services, such as Amazon’s EMR. The system is designed for two user groups: Image analysis specialists and end-users, e.g. pathologists. The image analysis specialists can script the system, use the Application Program Interface (API) in scripts, or write new modules for it. The end-users can work with the user interface which provides out-of-the-box functionality for many use-cases. We started building Orbit in 2009 and added successively more and more functionality to solve most of the image analysis problems related to the drug discovery domain.

We evaluate the application of this tool for three real world problems. Each of this application makes use of the map-reduce (17) based tile-processing approach of Orbit: The mapper step applies image analysis algorithms on tiles (which can be performed in parallel), and the reducer step joins the tile results. Orbit provides functionality for pixel classification, object segmentation and classification, and deep learning methods for complex heterogeneous object detection. These algorithms can be applied on different resolutions of the image which allows the coherence of different image context levels which is comparable to the way a human pathologist works: Use a low magnification lens to get an overview and analyse the context, then use a high magnification lens to analyse some details keeping the context around in mind.

Orbit Image Analysis is a versatile tile-processing engine for whole slide imaging. It is a modular system which can access image and meta data through several image providers, apply image analysis algorithms in map-reduce manner, and optionally use a scale-out infrastructure like Spark to execute the map-reduce tasks in a distributed computing environment. Out of the box it can connect to the open-source image server OMERO, or run in stand-alone mode to access local files from disk. Orbit has many machine-learning based algorithms built-in like pixel classification, object segmentation and object classification. A sophisticated deep learning segmentation method allows the detection of complex heterogeneous objects. Additional algorithms can be implemented easily and integrated as modules. Orbit comes with a script editor which allows the execution of Orbit scripts directly inside the user interface. This open-source program and some models like the glomeruli detection model are licenced under the GPLv3 and can be downloaded at http://www.orbit.bio.

## Results

Orbit has been applied on many real world use-cases in the drug discovery domain. Here we will describe the results of two of them briefly and describe a third one, which is based on deep learning, more in detail.

Idiopathic Pulmonary Fibrosis (IPF) is a life threatening disease where collagen deposits in lungs lead to a shortcoming in the oxygen-blood exchange. Lung tissue cuts have been stained with Masson’s Trichrome staining to mark the collagen on the slide images. Evidences for IPF are collagen deposits along alveoli walls and dense fibrotic areas (sometime called fibrotic foci).

For quantifying the collagen along the alveoli walls it is very important to exclude the collagen along the vessel borders, which is structural collagen not related to fibrosis. However, both, the structural and the alveoli wall collagen have the same staining colour (blue). In addition, one big challenge for the quantification is the very high staining variability between images which stems from different batches stained from different staining machines and people. The current gold-standard is a manually performed analysis of Masson’s Trichrome stained paraffin sections using a scoring system defined by Ashcroft et al. (19). Today, with the combination of WSI scanners and the availability of huge computation power, Orbit can be used to automatically and objectively quantify IPF images ((20); (21)).

The Orbit workflow is a combination of an exclusion model and a classification model. The exclusion model consists of four classes: Two exclusion classes (background and vessel border), and two inclusion classes (normal tissue and fibrotic area). Normally, an exclusion model is only used for a ROI definition, however here the ratio of the fibrotic area is used as a result.

A classification model which discriminates between background, tissue and collagen is applied inside the valid ROI defined by the exclusion model. The collagen along the alveoli walls and the collagen around vessels can be discriminated because the exclusion model is applied on low magnification (~5x), and the classification model on high magnification (20x or 40x) of the image. The ratio of alveoli wall collagen is used as a second result.

Both output parameters, fibrotic area and collagen, can be used individually or as an input for a linear SVM to predict the real IPF scores which has been manually defined by pathologists. Stritt et al. reported in (20) that using this method a correlation factor of 0.81 can be achieved. In (18) the authors show that using a larger training set a model can be created which outputs robust results for a large series of images gathered over a time period of five years with high staining variability.

Intraepidermal nerve fibre density (IENFD) is a biomarker for peripheral neuropathies, defined by the number of nerve fibres crossing the basement membrane between the dermis and epidermis per millimetre (22). The gold standard so far is the manual assessment of IENFD under a microscope using some guidelines (23). The aim of Seger et al. published in (21) is to standardize the quantification of IENFD in human skin biopsies and to therefore decrease the inter-rater variability.

Based on many skin biopsy images, an orbit model was created to recognize the nerve fibres. A special segmentation variation based on a high and low threshold edge detection algorithm, similar to the Hysteresis plugin of ImageJ, is used to segment and connect nerve fibres. Low threshold parts which are connected to high threshold parts are connected to the high threshold object, the other stand-alone low threshold parts are discarded. Technically this segmentation step is a mapper step. The reduce step connects partial nerve segments which are adjacent at tile borders.

The model including the threshold parameters is very robust (21) and can be applied to fluorescence and bright-field stained skin biopsies (Fig 3). It is embedded and freely available in the nerve fibre detection module in Orbit.

**Fig 1:**
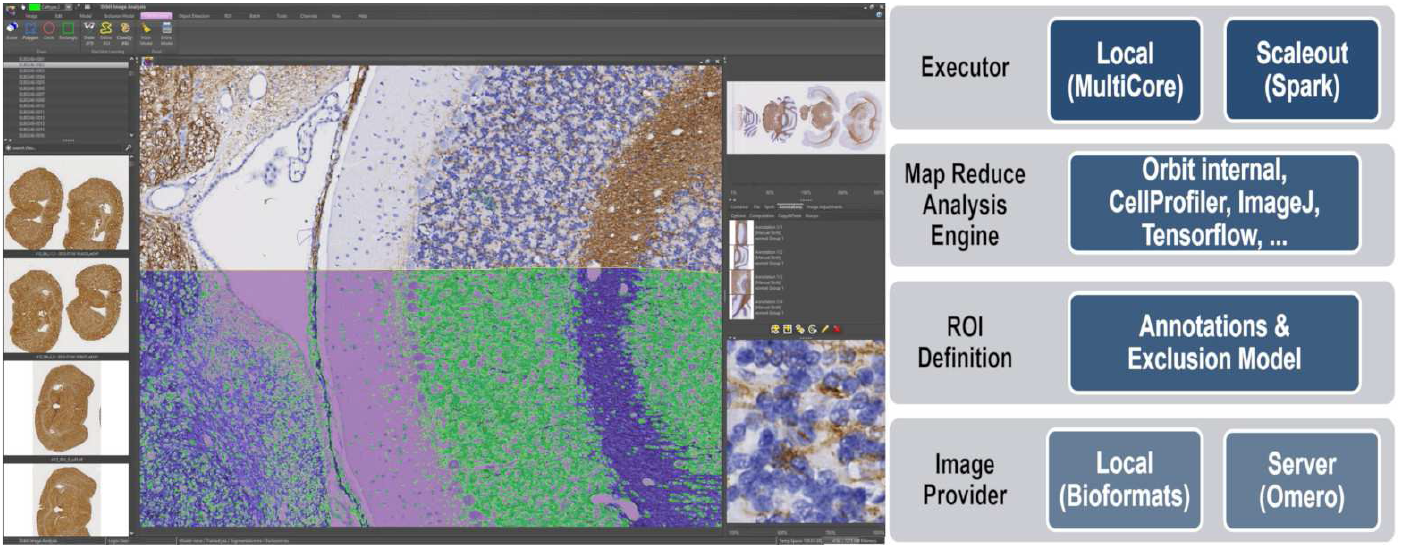
Overview of Orbit. Left: The Orbit user interface: Image browser on the left, tasks are on top grouped in tabs, properties and working results on the right. Image viewer in the centre. Right: The architecture of Orbit Image Analysis.

**Fig 2:**
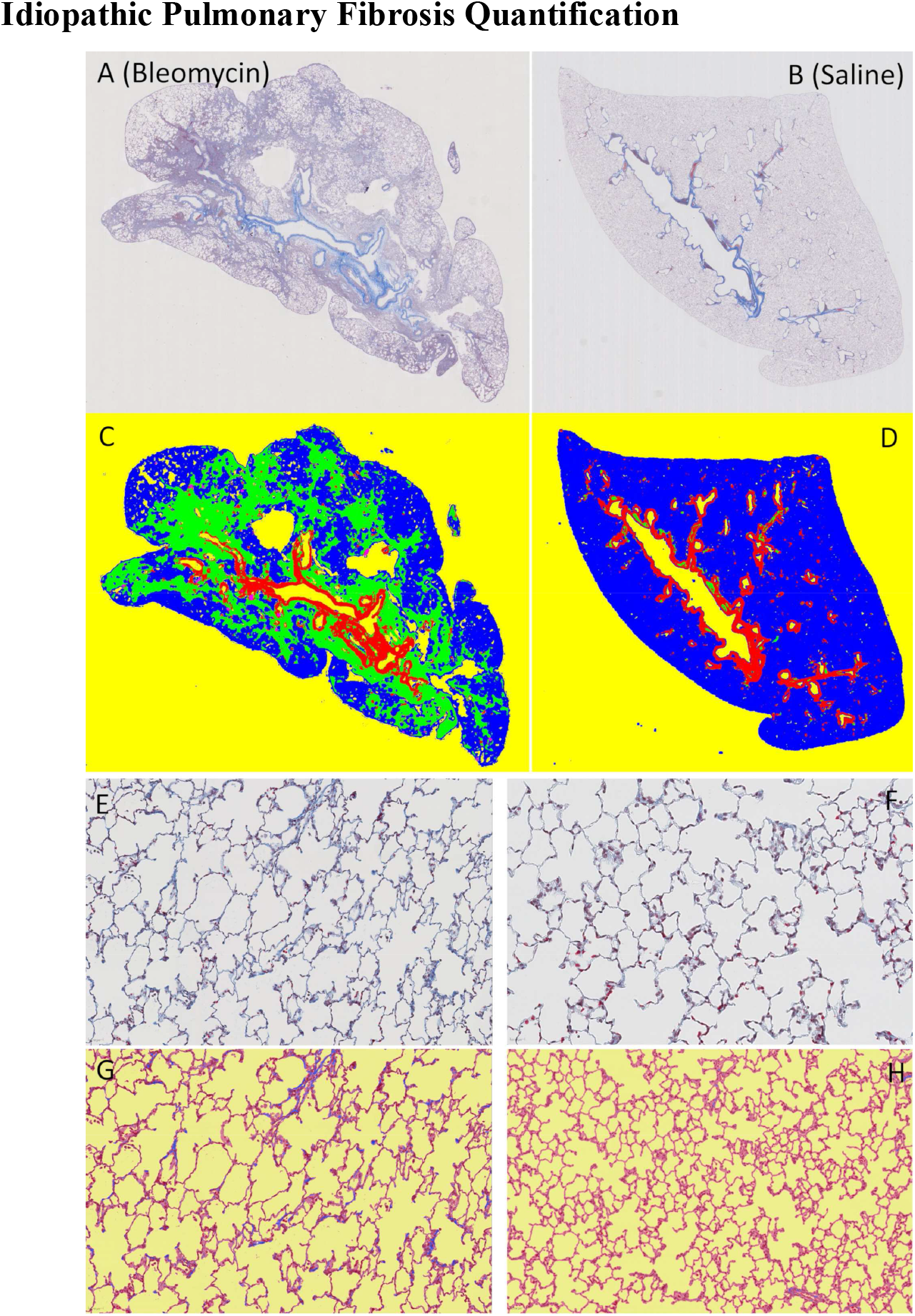
Idiopathic Pulmonary Fibrosis Quantification (IPF). IPF quantification: Treated (left) and control (right). (A,C) and (B,D) shows how the exclusion model works. (E,G) and (F,H) shows the finegrained collagen quantification.

**Fig 3:**
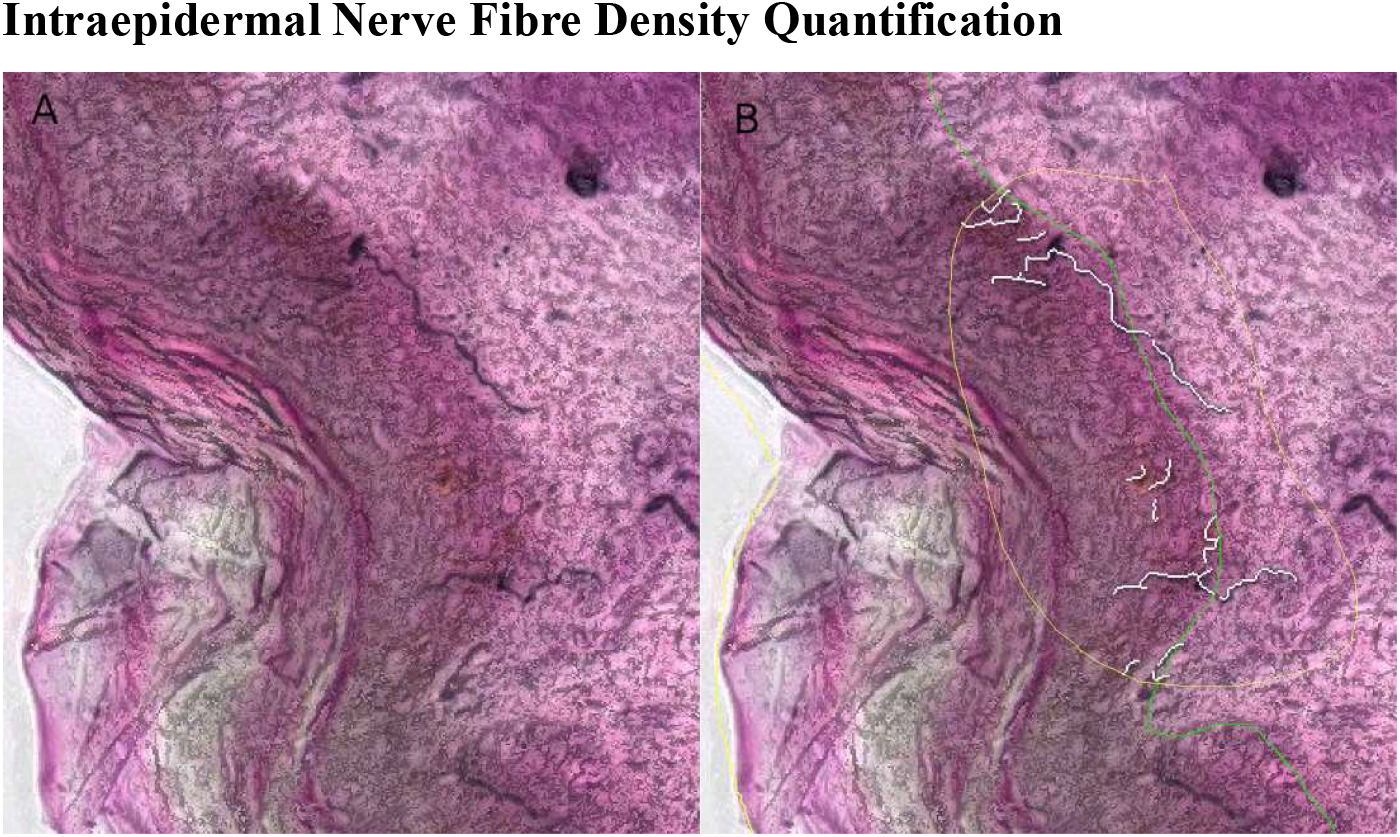
Intraepidermal nerve fibre detection. Nerve fibre image (A) and nerve detection (B) on a bright-field image. Only nerve fibres close to or crossing the manually annotated junction line are detected.

After automated nerve fibre detection all segments crossing the dermal–epidermal junction are randomized and presented to the user as a list. By clicking on an element the corresponding nerve fibre segment is centred on the screen and the user has the option to either skip it in case of false detection, or score the number of junction crossings by pressing a number key.

This method leads to a very robust and unbiased (randomized sections, each section presented only once) quantification which decreased the intra-rater variability.

Standard object detection methods allow the detection of homogeneous objects like cells, but fail on complex heterogeneous objects where a larger context plays an important role. A good example for this is glomeruli detection, which is important to quantify various kidney diseases. Glomeruli in kidney tissue are heterogeneous objects with a high variability. For detecting these objects, we apply a deep learning segmentation approach based on a encoder-decoder convolutional neuronal network (CNN) structure. This approach is generic and can be used to detect other objects than glomeruli as well (14). We provide a pre-trained model for glomeruli detection which is ready to use for segmentation. It can be downloaded from the Orbit model zoo: http://www.orbit.bio/deep-learning-models/. For detecting other objects one can build a new training set based on manually annotated objects which then can be used to train a new segmentation model.

### Training

To build a new deep learning model from scratch the first step is to annotate a set of representative objects over several images using the annotation tools. For the glomeruli example we annotated 92 slide images resulting in around 21’000 annotations. This huge amount of annotations was necessary because the aim was to create a model which is able to cope with two species (mouse and rat) and several stainings: H&DAB, FastRed, PAS, and three variations of H&E. The ultimate goal is to detect glomeruli on any WSI. In addition to the object annotations we drew a ROI annotation to define the region which we considered for the annotations. This implies that outside the ROI there might be further objects which are not taken into account.

**Fig 5:**
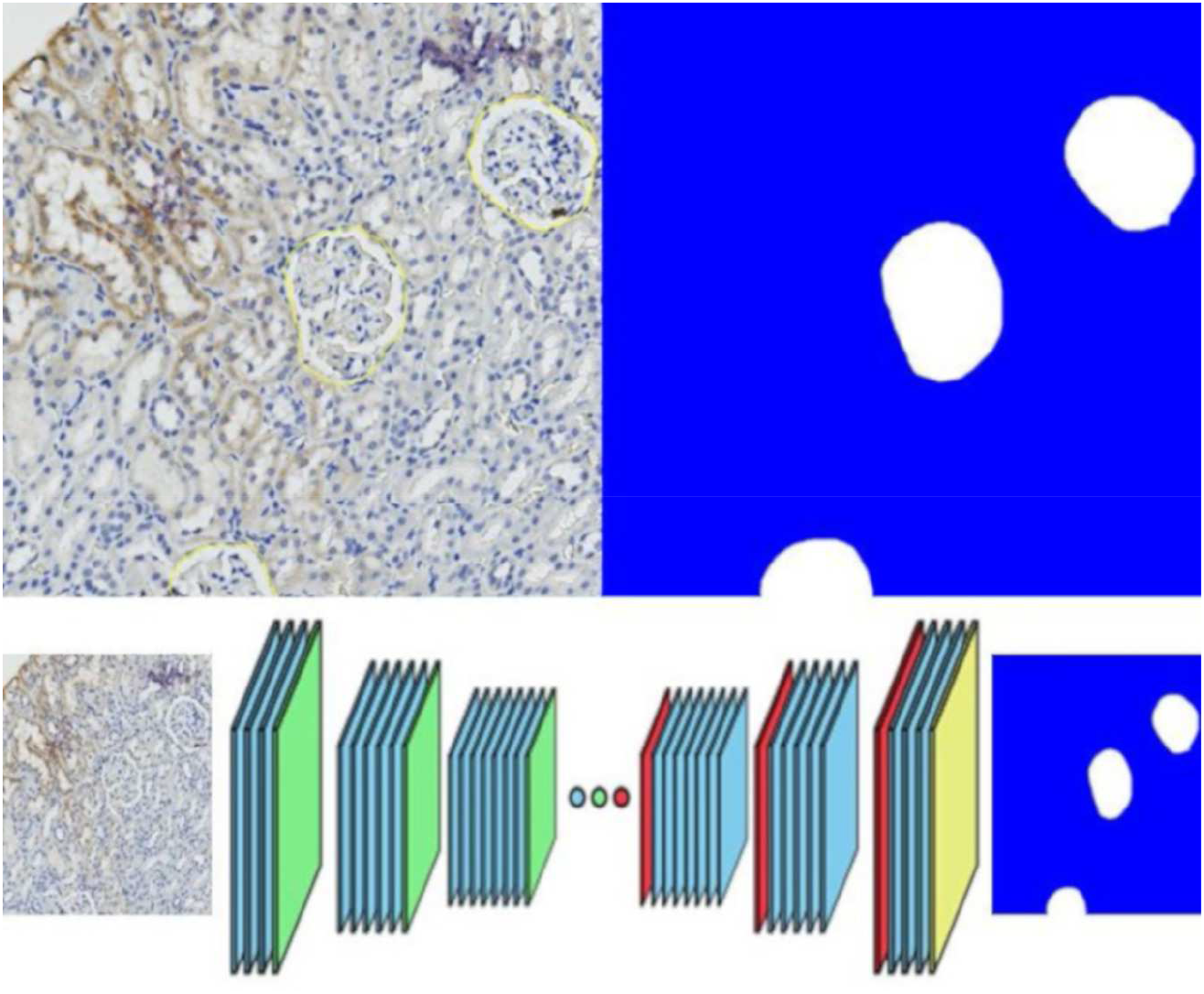
Deep-Learning object detection. Top: Annotated objects on left side, generated segmentation map which is used as training input on right side. Bottom: The ResNet-101 based encoder-decoder network takes a tile image as input and outputs the tile mask.

Orbit generates 512×512 pixel sized tile masks which encodes objects and background out of the manually drawn annotations [Fig 5]. These mask act as training set for the actual training of the deep learning model. We provide a Python script for training which has to be executed outside of Orbit, ideally on a machine with a GPU (http://www.orbit.bio/deep-learning-object-segmentation/). The script uses TensorFlow as a deep learning framework and runs out of the box, only the file path of the tile masks has to be defined. As a base CNN we use a ResNet-101 network structure (24) pre-trained on the MSCOCO dataset (25). The input tensor is a tile image and the output (target value) is its corresponding tile mask. The encoder-decoder network structure of it is visualized in Fig 5.

### Segmentation

Once the model is created, or a pre-trained model downloaded, objects on images can be segmented. As a first step the CNN model is used to predict the tile masks. We observed that the model detects objects very well if they are located in the centre of the tile, but often only partially when they are located close to the tile border. Fortunately, the false-positive pixel-rate in the tile mask is close to zero, meaning that if the model detects a pixel as inside an object it really is - it just misses some. This allows a segmentation refinement step: Arbitrary translations are applied to the tile, for each variant the tile mask is predicted, and the refined result mask is the union of all variant masks with respect to their inverse translation. For the glomeruli detection, four translations have been applied: Half the tile size up, down, left and right. Based on the tile mask the object segmentation is performed using a pre-defined segmentation model. Another refinement is applied to cope with tile-crossing objects: Each segmented object is centred, a virtual tile around the centre of the object is extracted and used as input for another segmentation step. The idea is that partially segmented objects are detected completely if they are located at the centre of the tile. All segmented objects are stored in an object list. A post-processing step eliminates duplicated objects from that list by only keeping the once which cover others.

### Evaluation

The complete dataset consists of 90 whole slide kidney images from rat and mouse samples and several stainings. Fig 4 shows the huge variability between the stainings. For this evaluation the data set was divided into 60 training and 30 test images. Each slide image contains around 300 glomeruli objects. For each group (species x staining) the test samples were independent but randomly selected to ensure a good stratification. The training set was virtually enhanced with 90- and 180-degree rotation images in order to achieve rotation invariance. The complete training set including the rotations contains 87213 images and masks, each 512×512 pixels. The training of the network has been performed on a high-end gaming GPU and lasted 4 days for 410’000 training iterations with a batch size of 2.

**Fig 4:**
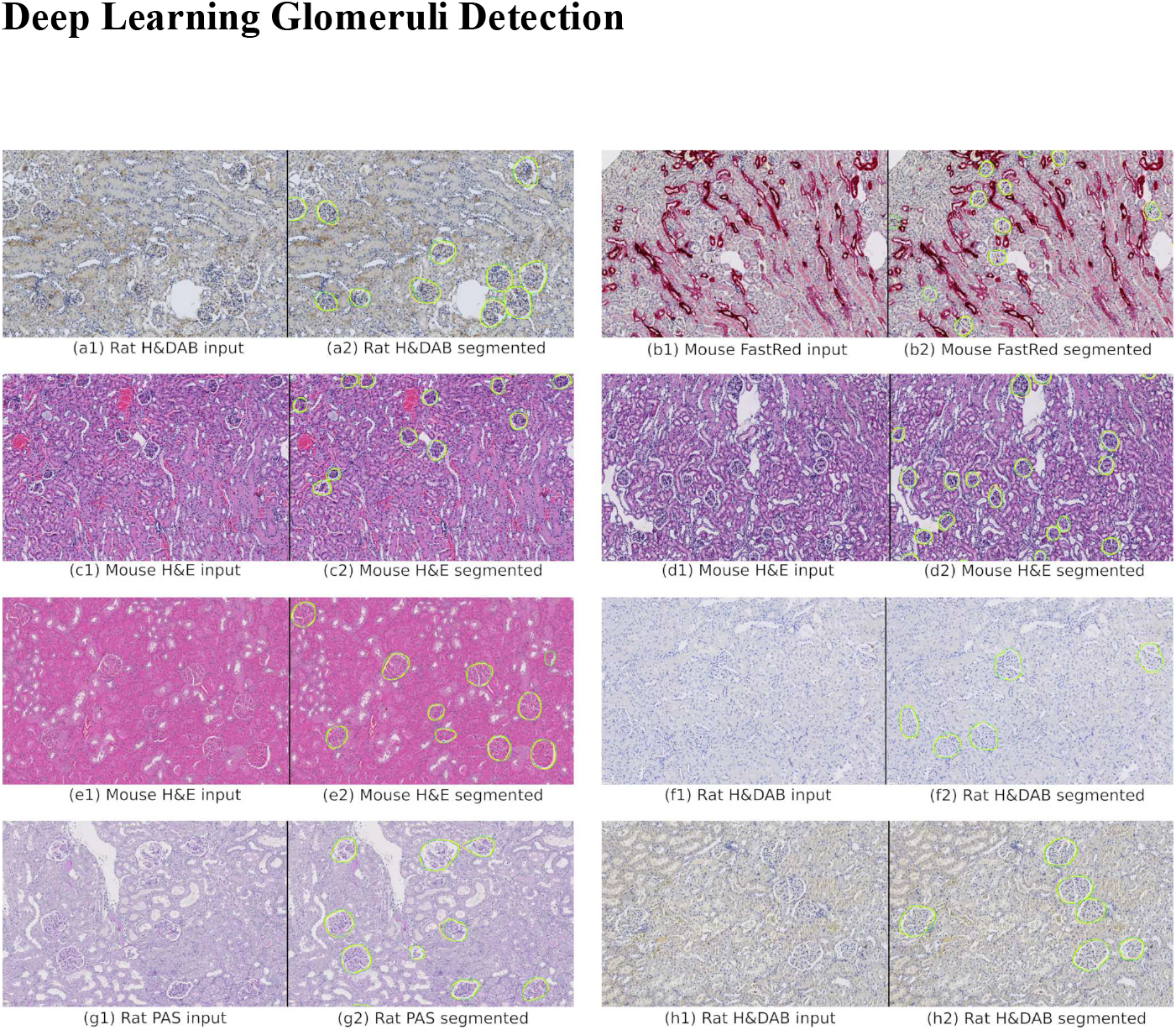
Glomeruli detection on kidney slides. Manual annotated glomeruli outlines (ground truth) in yellow, detected glomeruli outlines in green.

The evaluation of the test set is based on the Dice index (26) which scores the overlap between pixel in the rage of [0,1] where 1 means a complete overlap. For each ground truth object, the closest segmented object is taken into account and vice versa. The sum of the two overlap ratios is divided by two. We use the dice object index as defined in (27) to score a complete image by using the Dice index for all objects, weighted by its object size.

Table 1 shows the Dice object index results per group and total. The scores show a good segmentation performance over a set of 30 whole slide images, each containing around 300 glomeruli. This numeric evaluation has been confirmed by a subjective visual evaluation as is it exemplary shown in Fig 4.

**Table 1:** Evaluation results of the glomeruli segmentation for each group and total. Samples of each staining type are illustrated in Fig 4.

| Type | Mean Dice Obj. Index ± SD | #Images |
| --- | --- | --- |
| H&DAB Rat 1 | 0.8299±0.0194 | 4 |
| H&DAB Rat 2 | 0.8975±0.0100 | 4 |
| H&DAB Rat 3 | 0.8837±0.0245 | 4 |
| FastRed Mouse | 0.8225±0.0319 | 2 |
| PAS Rat | 0.9092±0.0129 | 4 |
| H&E Mouse 1 | 0.9030±0.0123 | 4 |
| H&E Mouse 2 | 0.8807±0.0203 | 4 |
| H&E Mouse 3 | 0.8768±0.0135 | 4 |
| Total | 0.8754±0.0181 | 30 |

Note that the ground truth is defined by manual annotations which might not be exact or miss some glomeruli. Indeed, a visual inspection could show that the automated detection found some new glomeruli which had not been annotated. Interestingly the models work very well over a high staining variability, e.g. H&E images differs a lot from FastRed and H&DAB images, but the model is able to detect glomeruli in all groups.

## Discussion

We showed the successful application of Orbit within three completely different image analysis domains: The lung fibrosis quantification is a structure discrimination task which demonstrates the need of combining several magnification levels of an WSI.

The intraepidermal nerve fibre density quantification showed the challenging task of segmentation across image tiles and combines it with semi-automated analysis methods: The computer presents segments to the user, and the user manually applies a grade. This is only possible if the analysis engine and the user interface smoothly work together.

The third application demonstrated the integration of a complex deep-learning model to identify glomeruli objects in kidney. This demonstrates two powerful abilities of Orbit: The integration of external tools, and the combination of the partial (tile-based) analysis results. Here we used TensorFlow to train the deep learning model and also as inference engine. The Orbit tile-processing engine then combined the partial results per tile. Only with that approach it is possible to detect glomeruli across tile borders.

We believe that the generic tile-processing engine of Orbit, combined with sophisticated machine-learning algorithms unveils the intrinsic power of WSI in digital pathology. It allows the integration of existing algorithms made for smaller images and enables the usage of them for very big images. The scale-out mechanisms, e.g. using Spark, allows to scale-up, and process hundreds of WSI in parallel. The scripting functionality facilitates the users to work in an agile process. The combination of all these aspects leads to an automatized and objective WSI quantification process.

## Methods

### Architecture

Orbit Image Analysis is a modular system which allows to exchange several components. Fig 1 shows an overview about the architecture.

#### Image Provider

Image providers provide tile-based image data, provide search functionality and read and store meta data like annotations and models. Out of the box Orbit comes with a local image provider which reads images from disk in standalone mode. The *BioFormats* library from the Open Microscopy Environment (https://www.openmicroscopy.org/) is used to read many proprietary WSI file formats. A unique ID based on the md5 hash of the image data is assigned to each image. Meta data is stored in a SQL-Lite database on local disk. In addition the image provider OMERO can be used to connect to the open-source image server OMERO (28). This image provider makes use of the OMERO-API to read tile-based image data and also to store meta-data. The authentication mechanisms and access rights of OMERO are used. This allows a collaborative workflow in a distributed environment, e.g. biologists create annotations, image analysis experts run analysis algorithms within these annotations. Other image providers can be integrated easily by implementing an *IImageProvider* interface so that other imaging infrastructures can be used as data source.

#### ROI Definition

For WSI the region of interest (ROI) definition is very important. This defines e.g. where the tissue on the slide is located, or even more fine-grained, which parts of a tissue should be analysed. For this, Orbit provides two ROI definition modules. First, manual annotations can be drawn using the annotation tool panel. Several tools including a ‘magnetic lasso’ tool can be used to define a combination of ROIs, exclusion parts and explicit inclusion parts (areas which will be included even if excluded by other rules like exclusion annotations). Second, a so-called exclusion model can be trained. This is a pixel-based machine-learning model trained on a lower resolution of the image to define inclusion and exclusion areas. This automated approach can be combined with the manual annotations, which have priority.

#### Map-Reduce Paradigm

Image processing is performed in map-reduce manner. The mapper step is performed on tile level; the reducer step aggregates all mapper results. This allows arbitrary algorithms, e.g. ImageJ (29), Fiji (30), or Cell Profiler (31), to run on small in-memory image tiles, even if the algorithm is not designed for WSI. A good example here is cell count: The mapper step counts cells in each tile (e.g. 512×512 pixel), the reducer step is simply a sum function. The advantage of this process is that the mapper steps can be executed independently from each other, which allows it to be executed multi-threaded on a local computer or distributed, e.g. using a Spark environment. Orbit comes with build-in algorithms like a sophisticated structure-based pixel-classification, object segmentation and object classification. In addition, other algorithms or tools like Cell Profiler, ImageJ Macros, or Deep Learning segmentation using TensorFlow (32), can be used in the mapper step.

A generic map-reduce framework has been developed which allows to implement mapper and reducer steps and then execute these steps using different executors. Orbit comes with a local executor which executes map-reduce tasks multithreaded on the local computer. In addition, a Spark executor can be used to execute tasks on a Spark cluster, if available. This scale-out functionality allows the use of on-premise clusters with a Spark installation, or the use of cloud-services, such as Amazon’s EMR. Further executors can be developed easily by implementing the *IMapReduceExecutor* interface.

### User Interface

The user interface enables easy model and annotation creation and can be used as an image viewer for WSI.

On the left side panel datasets and images can be searched and selected. The image provider defines the hierarchical structure, e.g. in OMERO images are arranged by groups, projects and datasets. A search field allows a quick search by filename.

The top tabs show tasks like browsing and saving images and models, image analysis tasks (violet tabs), batch processing, plugin extensions (tools) and a versatile Groovy script editor. Each task is organized in several ribbon buttons which usually should be applied in the order from left to right. The ribbons with bold font are mandatory steps, the others optional. A mouse-over help tooltip is displayed for each button which should limit the need of an additional documentation. In addition a handbook and tutorials are available under https://www.orbit.bio/help/.

The right panel shows working elements, such as annotations and models, and image specific tools such as image adjustments and meta data. For instance, the annotation tab provides functionality to draw ROIs, which then optionally are stored in the database.

In the centre pane images and mark-up are rendered. The Orbit render engine supports standard RGB (bright-field) and multi-channel (fluorescence) images. In the latter case each channel is assigned a hue value, which can be modified by the user in the *Adjustments* panel. The render engine works multi-threaded, so that several tiles of the image are rendered off-screen in parallel and then displayed when the render process has finished. Orbit makes use of image pyramids for a fast rendering of big images at low zoom levels. Several overlays like a classification mask, segmented objects or manual annotations are rendered on top of the image.

**Fig 6:**
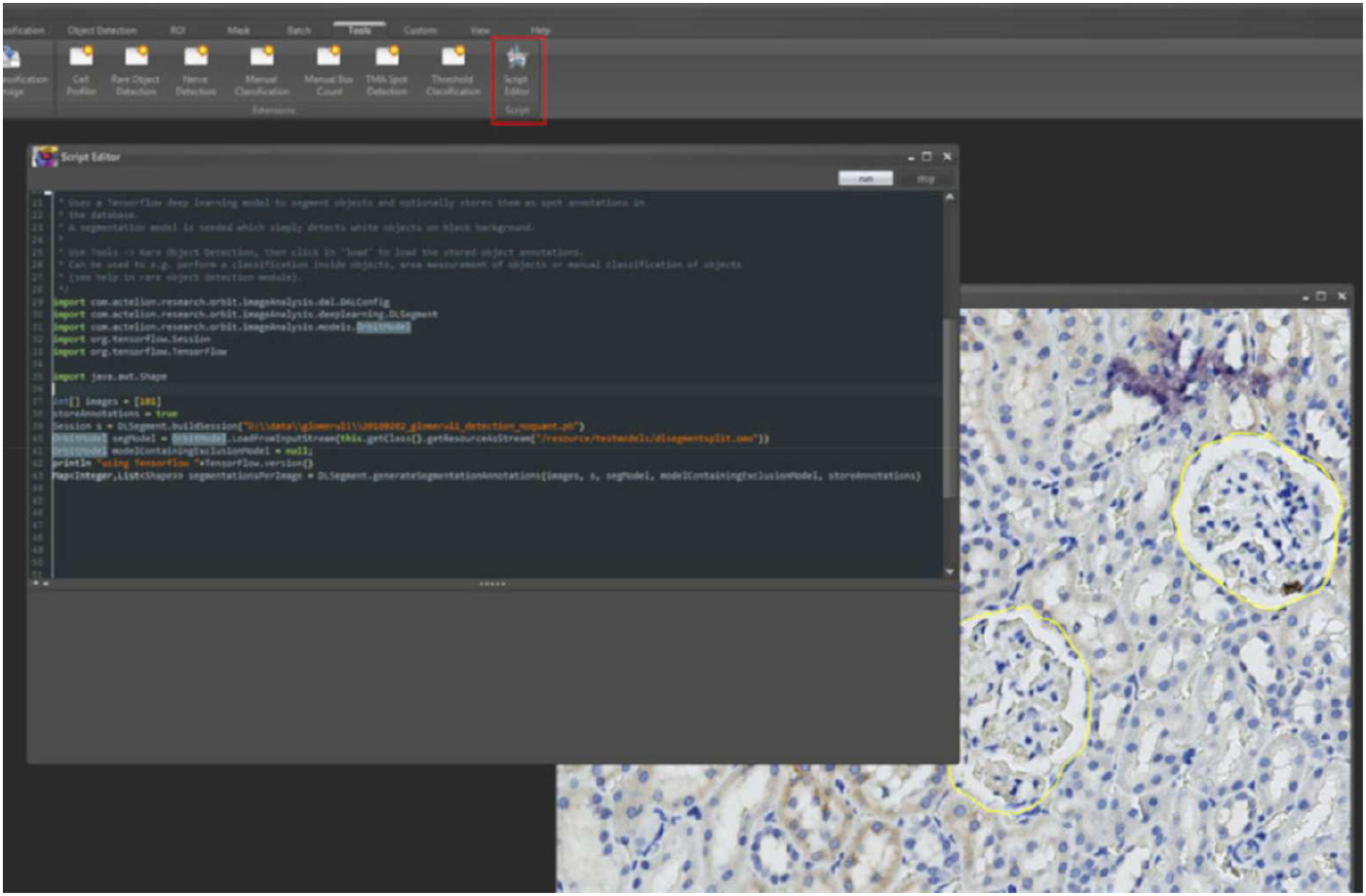
Integrated script editor. The integrated script editor allows the execution of Groovy scripts which can access the Orbit API. Results can be visualized as mark-up on open image frames.

#### Script Editor

An integrated Groovy editor enables the execution of Groovy-Scripts directly in Orbit. The advantage of the integrated editor is that the code execution runs within the same Java Virtual Machine (JVM) and thus can access all active objects and has the full Orbit plus dependencies class-path available. The main controller class *OrbitImageAnalysis* is a singleton and can be used to easily access all open images and models, read und process image data and even write back objects, e.g. polygons or an overlay map.

Scripting allows the automated execution of manually performed steps in the user-interface, and in addition perform complex analysis steps which are not integrated into the user-interface. Orbit provides sophisticated helper methods in the *OrbitHelper* class to handle ROIs, classify pixels, segment objects or classify objects. Image data can be requested to execute arbitrary algorithms on tile data, e.g. run an ImageJ macro on it.

The benefit of the script editor is that the workload of image analysis specialists and end-users can be split: The more programming oriented image analysis specialists can write parameterized scripts so that the end-users can adapt some parameters (e.g. a threshold), even without deep knowledge in scripting, run the script, validate the results, and eventually adjust the parameters. The validation can even be done visually if the script writes segmentation objects or a classification map back to the image.

For headless script execution the *GroovyExecutor* class takes a URL of a Groovy-Script as argument and executes the Groovy code. This allows to run Orbit on a server and to submit a script which is located in e.g. a source control system. This is supplementary to the map-reduce principle, actually a script itself can start map-reduce processes.

### Quantification Methods

Orbit comes with sophisticated build-in image analysis algorithms. The main principle is a machine-learning based pixel classification which is used for many other algorithms like object segmentation. All algorithms work tile-based, which means tile by tile is processed. However, only the tile index is specified in each step, it is up to the algorithm if only specific tile-data is requested or a larger context, e.g. surrounding tiles. Algorithms can also be performed on different image resolutions, for example an exclusion map is performed on a very low resolution to get a better overview of the whole slide.

#### Pixel Classification

One of the basic principles in Orbit is to classify pixels. Therefore, the user defines a set of classes, e.g. ‘background’, ‘staining 1’, ‘staining 2’ and assigns each class a colour. One important parameter to set is the so-called structure size. This parameter defines the size of the surrounding context taken into account for each pixel. For each class the user draws several shapes using a polygon, rectangle or circle tool. The union of all pixels inside these training-shapes define the training set. Several images can be used to define the training set with the goal to cover the whole variability of the tissue structures. For each pixel inside the training set a feature extraction is performed, which takes not just the pixel itself, but some area around each pixel defined by the structure size, into account. Based on this data the following features are computed: Min, max, mean, standard-deviation, edge-factor, and the middle pixel intensity per channel. For example, the edge-factor for pixel *p* characterizes the edginess and is defined as

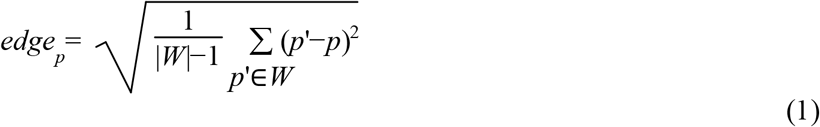

*W* defines the set of pixels inside the structure-size window around pixel *p*.

These features are used as input for a Support Vector Machine (SVM) which is used to train a statistical model for pixel classification. That way the user does not have to specify any thresholds, the tissue class discrimination is done simply by drawing shapes on representative regions. The structure-size here is very important: Smaller values discriminate more fine-grained, higher values take more context into account.

#### Object Segmentation

A classification model is used to discriminate foreground and background pixels. The segmentation task groups connecting pixels to an object and defines a shape which surrounds it. Object segmentation in the biology domain is mainly used to segment cells. Here the main challenge, especially in tissue, is that cells might overlap. Orbit provides a standard watershed algorithm (33) in combination with erosion and dilation steps which solves most of the problems. For challenging problems like dense cell clusters a Mumford-Shah functional (34) based segmentation algorithm can be used.

Two-level segmentation performs a second segmentation inside segmented objects. For this a second classification model is build and applied within the detected objects of the primary segmentation model. This is very useful for e.g. mRNA detection which is small spots inside cells.

By default, Orbit simply outputs the number of segmented objects, however, in the segmentation settings the *Features* option can be activated. This enables the feature computation per segmented object which computes several parameters like shape descriptors, texture descriptors, location information, and outputs it per segmented object.

**Fig 7:**
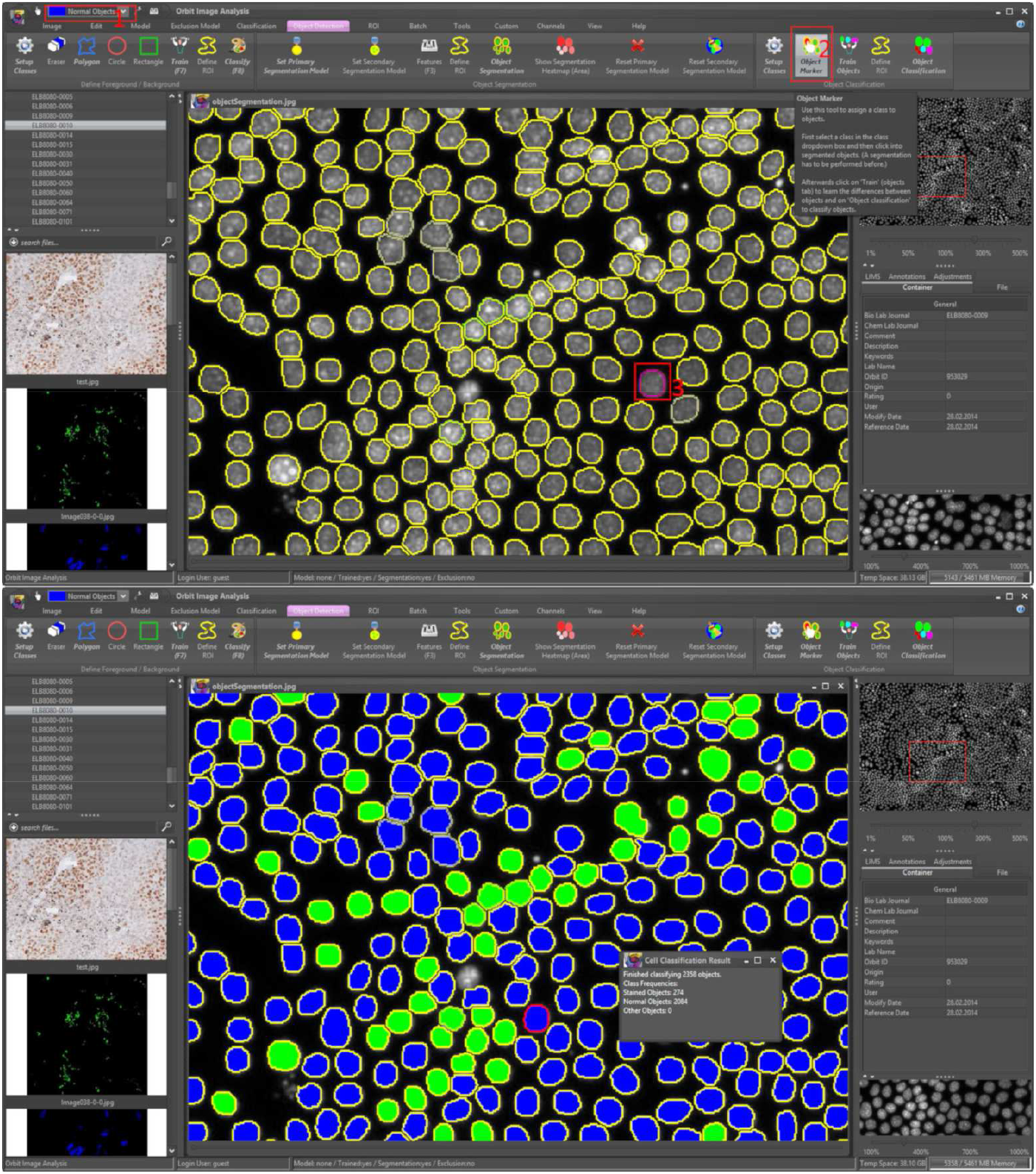
Object classification. Upper image: After an object segmentation the user can define object classes (1), user the object marker tool (2) and mark segmented objects (3). Orbit computer features per object and uses a SVM to classify the objects. Lower image: Results of the object classification.

#### Object Classification

A machine-learning based object classification can be applied after the segmentation. The idea is that the user can mark segmented objects and assign them to specific classes - without thinking about how to describe differences between the object classes. To achieve that, Orbit provides an *object marker* tool which can be used after a segmentation step. For each class, the user marks several objects which defines the training set. Orbit computes internally the object features per segmented shape which will be used to train a SVM. Based on this statistical model all segmented objects are labelled. The object classification can be seen as an additional step to the segmentation. Sometimes this is very useful to discriminate objects, sometimes the output of the object feature output in combination with an external data analysis is more appropriate.

#### Region of interest

The region of interest (ROI) is the area in which other algorithms are applied, e.g. an object segmentation only segments objects within the valid ROI and skips the other part of a WSI. Usually the ROI defines the tissue and excludes the background, but it can also exclude unwanted parts like holes or wrinkles.

#### Annotations

Manual annotations can be drawn in the annotation panel using annotation tools like a pencil or a magnetic lasso. By default, an annotation can have a name and is just an informative annotation. However, annotations can be edited and given a special type. Annotations of type *ROI* define the region of interest (e.g. area outside ROIs is excluded). If multiple ROIs are created, the final ROI for the image is the union of all ROI annotations. *Exclusion* annotations define excluded areas (e.g. inside a ROI annotation), except another annotations of type *inclusion* exists. Inclusion annotations have the highest priority and overrule exclusion annotations. All created annotations are saved in the database and automatically loaded whenever an image is accessed.

#### Exclusion Models

Technically an exclusion model is a classification model on a low resolution (approx. one megapixel) with a high structure-size. For each class, the user can define whether it is an inclusion or exclusion class. The training is done the same way a standard classification model is trained but a special exclusion model training automatically uses a low resolution of the image and later maps it to the full resolution. Manual annotations (e.g. exclusion annotations) and exclusion models can be combined, so some parts of a WSI can be explicitly excluded, even if the exclusion model would include it.

#### Masks

The masking functionality in Orbit is a generic approach to define areas which are included and areas which are excluded from analysis. In contrast to the exclusion model and manual annotations, masks do not affect the measured size of a ROI.

A classification mask is a standard pixel classification model with additional flags for each class defining whether the class should be included or excluded. In contrast to an exclusion model the model usually us applied on a high image resolution. The classification of the pixels is performed on the fly tile by tile when main analysis on the particular tile is performed. One speciality here is, that the mask classification and the main analysis can be applied on different channels. This allows a channel colocalization quantification like it is often used in fluorescence imaging.

A segmentation mask is a segmentation model in which all pixels inside segmented objects are declared as active pixels used by further analysis. This allows a pixel classification inside segmented objects, or a nested segmentation.

#### Tools

Orbit provides many other tools out-of-the-box to cope with the most common image analysis problems. This includes among others a tool to run Cell Profiler pipelines within the ROI defined by Orbit and read back the cell segmentations, a nerve fibre detection tool, a tissue microarray (TMA) spot detection tool and a rare event detection tool. The latter is a very flexible tool which performs an object segmentation and presents all detected objects to the user who then can manually assign each object to a class. A report outputs the frequencies of the assigned classes.

In addition to the build-in tools, image analysis specialists can create custom tools which are instantiated via Java reflection and listed as ribbons in the *custom* tab. The developer has access to all open images, annotations and models in order to perform an analysis task.

## Acknowledgements

We would like to thank Shanon Seger for her work on the IPF and IENFD models, and Julia Marrie and Celine Runser for the scientific input, model evaluation and sample preparation. We would like to thank Thomas Sander for supporting this project and Aaron Hart for the critical reading of this paper. The authors thank the OME team for help with OMERO integration and Jason R. Swedlow for critical reading of the manuscript.

